# Sharing GWAS summary statistics results in more citations: evidence from the GWAS catalog

**DOI:** 10.1101/2022.09.27.509657

**Authors:** Guillermo Reales, Chris Wallace

**Affiliations:** Cambridge Institute of Therapeutic Immunology and Infectious Disease (CITIID), University of Cambridge, Cambridge, United Kingdom; Department of Medicine, University of Cambridge, Cambridge, United Kingdom; MRC Biostatistics Unit, University of Cambridge, Cambridge, United Kingdom

## Abstract

Genome-wide association studies (GWAS) have been a crucial tool in genomics and an example of applied reproducible science principles for almost two decades.^1^ Their output, summary statistics, are especially suited for sharing, which in turn enables new hypothesis testing and scientific discovery. However, GWAS summary statistics sharing rates have been historically low due to a lack of incentives and strong data sharing mandates, privacy concerns and standard guidelines.^2^ Albeit imperfect, citations are a key metric to evaluate the research impact. We hypothesised that data sharing might benefit authors through increased citation rates and investigated this using GWAS catalog^3^ data. We found that sharers get on average ~75% more citations, independently of journal of publication and impact factor, and that this effect is sustained over time. This work provides further incentivises authors to share their GWAS summary statistics in standard repositories, such as the GWAS catalog.

## Main text

In recent years, we have witnessed an increasing and solid push toward open science in the form of incentives for open-access publishing and data sharing across scientific fields, exemplified by Plan S (https://www.coalition-s.org/) and the rise of the FAIR (**F**indable, **A**ccessible, **I**nteroperable, and **R**eusable) principles, created as guidance for good data sharing practice to support data reusability.^4^ This effort comes from recognising that the accessibility and reuse of research data have a huge potential to boost scientific progress, especially given the vast amounts of data generated in genomics and biomedical fields.^5^

Human genomics pioneered the establishment of norms for data sharing, starting with the Human Genome Project, reflected in the “Bermuda principles” (https://web.ornl.gov/sci/techresources/Human_Genome/research/bermuda.shtml) and later expanded by the Fort Lauderdale agreement,^6^ which promoted the publication, sharing and maintenance of a community resource of genetic data, paving the way for successful multinational collaborative work.

Genome-wide association studies (GWAS) have been the workhorse of genomics for over a decade and are an example of reproducible science principles in practice due to the sharing of results and data.^1^ GWAS typical output, summary statistics (i.e. plain text files with the results of the per-SNP tests), are especially suited for sharing, as they are easily stored, alleviate privacy concerns posed by sharing individual data, and can be exploited by many bioinformatic techniques (eg. meta-analysis,^7^ Mendelian randomisation,^8^ linkage disequilibrium score regression,^9^ colocalisation,^10^ polygenic risk scores),^11^ thus enabling the reuse of existing data to explore new questions.

The NHGRI-EBI GWAS catalog^3^ is a publicly available and manually curated resource of human GWAS, which not only provides the most significant results and metadata of published GWAS but also offers structured and harmonised GWAS summary statistics associated with each study when available. However, there is still no agreement on GWAS summary statistic format, although efforts to develop one are being made^12^ or sharing policy, and recent work shows that most authors don’t share their GWAS data.^2^

Lack of data sharing is a common phenomenon across fields, and factors influencing data sharing have been investigated elsewhere (eg.^13,14^). Within GWAS, one particular challenge is participant privacy since individual-level genetic data is theoretically identifiable,^15,16^ and some possibility of identifiability exists even in summary statistics,^17^ although either would require someone to hold the genetic data on an individual already to identify them within a published study. Despite these concerns, in 2018, after considering all the risks and benefits, NIH supported the open sharing of summary-level GWAS data (https://grants.nih.gov/grants/guide/notice-files/NOT-OD-19-023.html).

There is still no definitive answer to which incentives would act to increase data sharing.^18^ We hypothesised that data sharing might benefit authors regarding citations upon data reuse. If this were so, it would provide an additional incentive, beyond good citizenship, for data sharing. We, therefore, used data from the GWAS catalog^3^ to explore the current sharing landscape of human GWAS summary statistics and to analyse the relationship between sharing and potential citations.

We collected sharing and citation information from 5756 studies with results published in the GWAS catalog.^3^ Roughly one in ten (604, 10.5%) had summary statistics available for download. The proportion of summary statistics-sharing studies has increased over the years, especially since 2015, but even in 2021, only 121 out of 578 studies (~21%) shared their summary statistics. (Fig. 1). Although we considered the GWAS catalog as the prime source of GWAS summary statistics, some datasets might be available elsewhere (e.g. authors’ or consortium’s websites or alternative repositories), making studies be mislabeled as non-sharers. To verify that our measure of sharing - whether the summary statistics were available in the GWAS catalog - was valid, we manually inspected a random sample of 353 manuscripts (out of 629) from two journals with high levels of GWAS publications, PLoS Genetics and Nature Genetics. We found that 324 (91.7%) did not provide full summary statistics or data was controlled-access, 5 (1.4%) claimed to provide access, but links were either broken or contained no data, and only 24 (6.8%) linked to full summary statistics in non-GWAS catalog websites (Table S1).

**Fig. 1.**
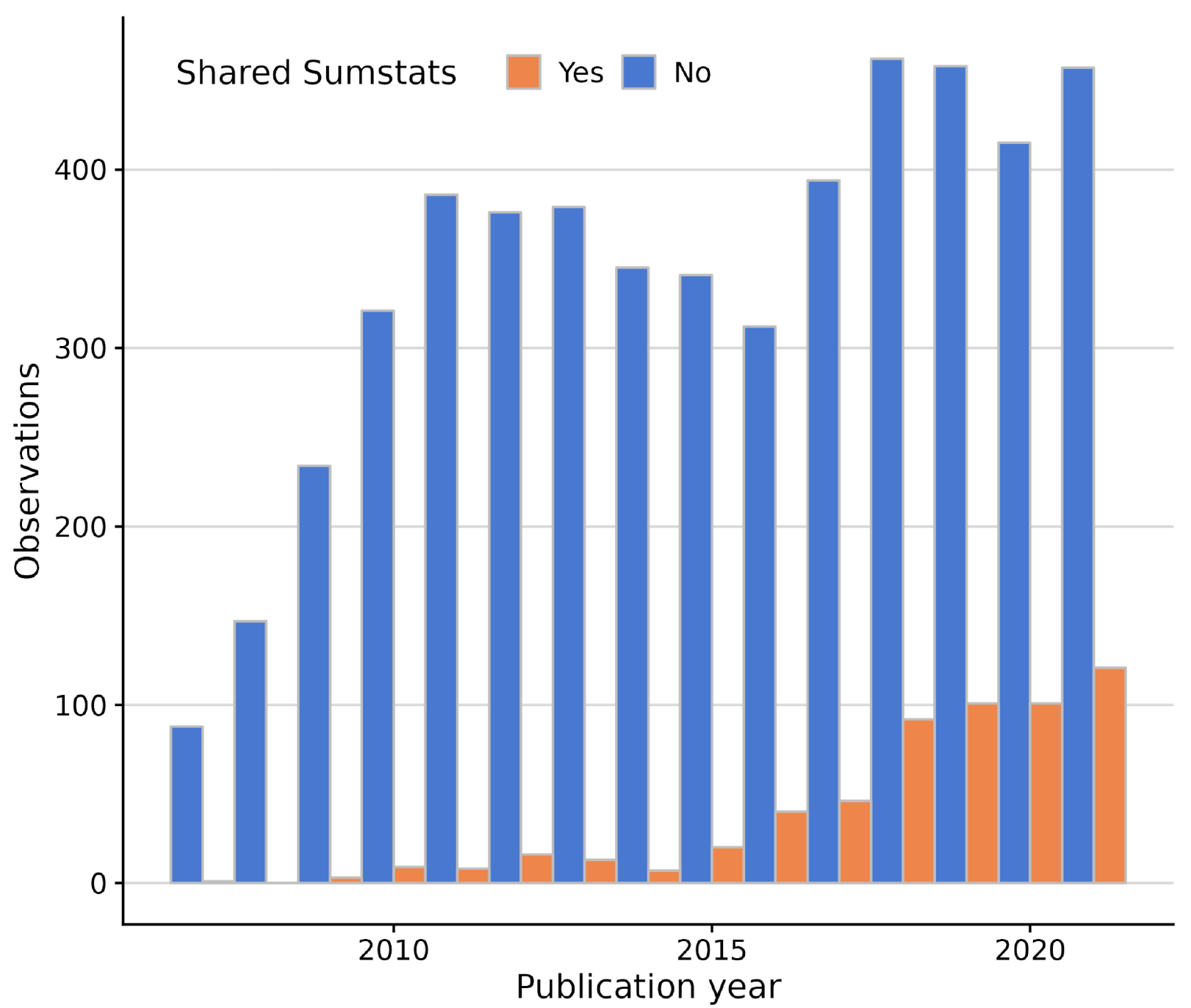
GWAS summary statistics sharing patterns in GWAS catalog, by year (2007-2021). Despite increased sharing from 2015 onwards, most GWAS studies do not share their summary statistics.

Satisfied that this was a valid measure, we used logistic regression to study which factors influence sharing in the GWAS catalog. According to the Bayesian Information Criterion (BIC), the optimal model included the year of publication, journal of publication and journal impact factor. Both year (OR = 1.4827 [1.4193 - 1.5489]) and journal impact factor (log(SJR) OR = 2.7820 [2.2242 - 3.4798]) have positive effects on sharing. Publishing in nine of the 20 journals that publish the most GWAS also significantly affected sharing (Table S2).

We decided to investigate the impact of sharing on a paper’s impact using the relative citation ratio (RCR), which compares the number of citations an article has to the average citation rates of the journals in its co-citation network.^19,20^ In the early years of GWAS, such articles appeared to outperform their co-citation network before a gradual decrease in the median score (towards RCR = 1), except for the most recent year, 2021. This bump may reflect incomplete data or a sudden behaviour change (Fig. 2a). As a broad pattern, studies that shared their summary statistics in the GWAS catalog had consistently higher RCR over the years than their non-sharing counterparts (Fig. 2b). Again, the data from 2021 appeared anomalous, with sharing papers showing only a weak advantage over non-sharing papers.

**Fig. 2.**
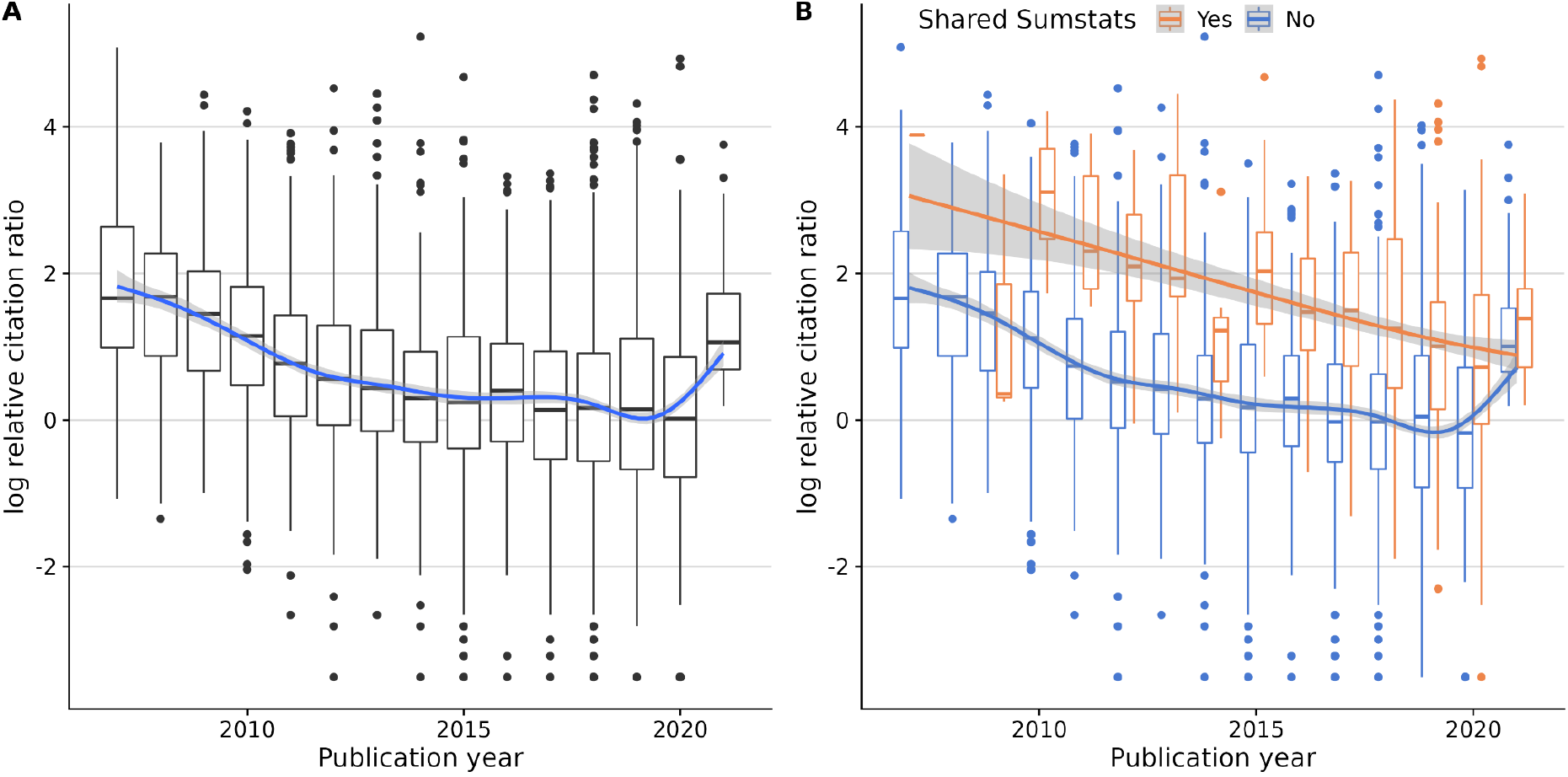
Citation patterns over time (2006 - 2021), measured in log relative citation ratio. **A.** All GWAS. **B.** Split by SumStat sharing status. Sharing studies are consistently more cited than non-sharing studies.

To try and understand the 2021 data, which had the shortest follow-up time by definition, we analysed the citation patterns of sharing and non-sharing studies over time by year of publication (Fig. 3). On average, GWAS citations rise quickly and stabilise around two years after publication. Then citations either stay stable or slowly decrease throughout the following years. However, summary statistics-sharing GWAS citation counts grow faster (Fig. 3a) and sustain higher mean citation counts, regardless of the year of publication (Fig. 3b). Given the citation advantage of sharing papers to non-sharing takes two or more years to accumulate, we decided to exclude the anomalous data points from 2021 because there had not been sufficient time for them to stabilise.

**Fig. 3.**
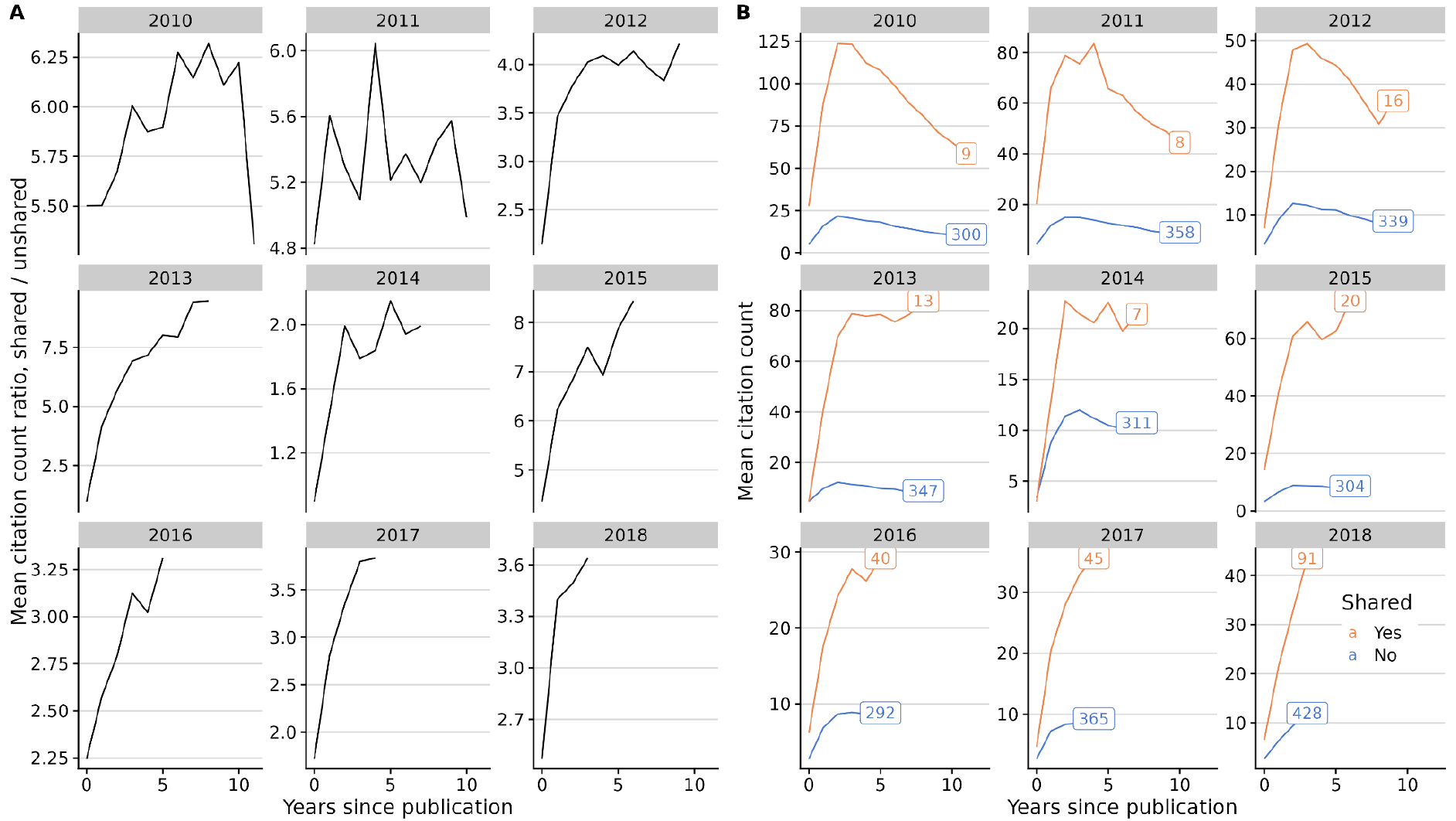
Mean citation count evolution after publication, by year of publication (2010 - 2018). Sharing studies get more citations from early on, then stabilising circa two years after publication **A.** Mean citation count ratio (shared/unshared). **B.** Sharing (orange) and non-sharing (blue) mean citation count. Text in squares indicates the number of studies in each category.

To analyse the effect of sharing on citations, we first built an optimal linear model of log(RCR) using all available covariates *except* sharing status according to the BIC. The selected model included the year of publication, journal of publication, journal impact factor, and the National Library of Medicine’s (NLM) “molecular/cellular”score.^19^ By adding a binary variable describing sharing practice, we concluded that sharing summary statistics has a positive effect on the RCR, providing ~75% more citations on average than non-sharing articles (RCR ratio = 1.7499 [1.5987 - 1.9154], P < 2e-16, Table S3).

Data sharing in the life sciences remains a controversial topic. We showed using GWAS catalog data that overall summary statistics sharing rates are low, although we see a remarkable increase in the past five years. Many factors not included in this work but analysed elsewhere,^21^ such as changes in scientific culture towards sharing, growing incentives from public and private funders, and varying privacy regulations across countries, along with technical difficulties, may influence sharing of GWAS summary statistics and other datasets. This may be further complicated by the multifactorial nature of data in many cases, the lack of clear definitions of what constitutes shared data, and the challenge of verifying the completeness of any dataset. Funders like Wellcome Trust, the NIH, and the ERC have mandated open-access publishing for articles, but strong mandates on data sharing are still generally lacking, and journal policies on existing data are not consistently enforced.^22^ Thus, while data sharing remains reliant on the goodwill and diligence of researchers, both the inertia to changing practice and the effort required may outweigh the limited incentives, leaving data unshared.

Citations are imperfect yet crucial metrics for evaluating research impact, which affects hiring decisions and career prospects. We hypothesised that sharing GWAS summary statistics may positively affect citations by allowing other scientists to conduct research using shared data and, in turn, cite the original research. Indeed, we observed a consistent pattern of increased citation rates over time, and by using linear models, we estimated that sharing increased citation rates by 75% on average, an estimate similar to the 68% increase in citations found in a study of microarray data sharing more than 15 years ago (Piwowar et al., 2007), and much higher than the 25% increase predicted in papers linking to more general biological data repositories.^24^

Our analysis of 353 GWAS papers that did not use the GWAS catalog revealed that most studies did not share data at all or shared either restricted access and/or incomplete data (e.g. only top significant hits), which hampers reuse. Only 24 articles shared full summary statistics without controlled access or request requirements using alternative repositories, and five provided links that did not work anymore, highlighting that the GWAS catalog has become the de facto standard for unrestricted summary statistic sharing as well as a reliable, future-proof data storage platform. Therefore, we encourage authors to use standard repositories like GWAS catalog whenever possible.

While appreciating the issue’s complexity, we support the implementation of more data-sharing mandates and recognition-based incentives, such as alternative metrics to promote data-sharing work, independent of journal of publication, as well as the inclusion of data generation and stewardship on researchers’ CVs.^25,26^ We also agree with other authors that the nature of increasingly large and more complex data sets will require improved training on data stewardship.^13^

We consider that the strongest incentive for scientists to share data is good citizenship because data sharing increases the ability of all of us to make discoveries through meta-analysis or integrative studies, thus accelerating scientific knowledge. However, and despite the observed recent trend changes, that incentive alone is clearly insufficient because papers sharing data remain a minority. We hope the robust evidence here that data sharing can increase citations independent of the journal of publication will provide further incentives and that we will see sharing of summary statistics continue to increase in the coming years.

## Methods

The GWAS catalog^3^ is an established and high-quality repository of curated human GWAS results, providing easy access to summary statistics made public by authors (via curator inclusion or author submission). Its large coverage (400,000+ associations from 5,690 publications as of May 2022) and its easy-to-access statistics make it an ideal reference database for our analyses. Hence, we downloaded the full list of studies and available summary statistics in GWAS catalog on 26th May 2022.

We fetched citation information for each study from NIH’s database using iCiteR v0.2.1,^27^ a wrapper for NIH’s iCite API.^28^ To quantify citations, here we focused on relative citation ratio (RCR), an improved metric to quantify the influence of a research article by using co-citation networks to field-normalise the number of citations.^20^ We also used iCiteR to retrieve the number of citations each study received each year.

Despite not being an appropriate indicator for the individual quality of a given paper, journal impact factor can affect citations via journal visibility and prestige. We retrieved 2021 SJR (SCimago Journal Rank) scores to assess overall journal prestige.^29,30^ There were 723 journals in our dataset, from which 696 had SJR data available. Those 27 without SJR data were either too new to have scores (eg. Nature Aging, EISSN: 2662-8465), changed names (eg. BMC Genomic data, ISSN: 2730-6844, previously known as BMC Genetics), or they were discontinued in Scopus (eg. Annals of Translational Medicine, ISSN: 2305-5847).

We additionally considered factors for the 20 journals with the most published GWAS to allow for additional variation between journals, pooling the rest as a reference category. The top 20 journals are: Am J Hum Genet, Am J Med Genet B Neuropsychiatr Genet, Ann Rheum Dis, Diabetes, Eur J Hum Genet, Front Genet, Gastroenterology, Hum Mol Genet, J Allergy Clin Immunol, J Hum Genet, J Med Genet, Mol Psychiatry, Nat Commun, Nat Genet, Nature, PLoS Genet, PLoS One, Proc Natl Acad Sci U S A, Sci Rep, Transl Psychiatry (Table S4).

We used the glm function in R 4.1.2^31^ to fit (1) a set of logistic models to explore the effects of time, journal of publication and other available factors on sharing, and (2) a set of linear models to explore the effect of sharing and other available factors on RCR. We chose to include all datasets published between 2007 and 2021 only, with 2007 being the first year with a shared summary statistics dataset and 2021 the last complete calendar year.

iCite tool uses Medical Subject Headings (MeSH) terms in articles’ text to predict the potential for translation of research.^19^ The tool provides scores that represent the proportion of terms that can be classified within three overarching branches of the MeSH ontology: Human, Animal, and Molecular/Cellular.

For each set of models, we sequentially added and removed predictors, using the BIC to choose the optimal model. For (1), this procedure selected the logistic model:

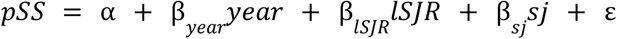

where *pSS* stands for public summary statistics dataset available, encoded as [0, 1], *year* is the year of online publication [2007 – 2020], *ISJR* is the logarithm of the SJR score, log(SJR), and *sj* stands for individual journal category, either one of the top 20, or other.

For (2), we selected covariates excluding *pSS* which produced the baseline linear model

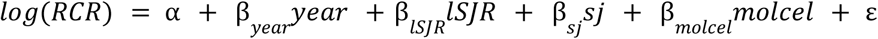

where *molcel* corresponds to the NLM molecular/cellular score, which showed to contribute to model fit, which we compared to

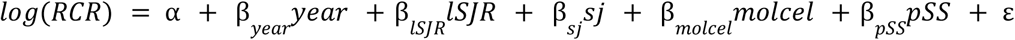

to quantify the effect of sharing on log(RCR).

While we expect manually-curated GWAS catalog to contain most publicly available summary statistics datasets, authors can choose to share their data on a different platform (eg. their own or consortium’s website, Dryad, or GWAS archive), posing a potential bias in our analysis. To explore this scenario, we selected random 50% of studies labelled as non-sharers in two of the journals with most published GWAS (PLoS Genetics (100 studies) and Nature Genetics (253 studies)) and manually checked whether their summary statistics were listed in the manuscript as freely available elsewhere and whether the statistics still resided at any such URL.

## Data availability

All source data used are available at the accompanying GitHub repository: https://github.com/GRealesM/gwas-sharing.

## Code availability

All code used in this work is available at https://github.com/GRealesM/gwas-sharing.

## Acknowledgements

C.W. and G.R. are funded by the Wellcome Trust (WT220788). CW is funded by the Medical Research Council (MRC; MC UU 00002/4) and supported by the NIHR Cambridge BRC (BRC-1215-20014). The views expressed are those of the author(s) and not necessarily those of the NHS, the NIHR or the Department of Health and Social Care. The authors thank all GWAS researchers who pro-actively share their full summary statistics on GWAS catalog or elsewhere.

This research was funded in whole, or in part, by the Wellcome Trust WT220788. For the purpose of Open Access, the author has applied a CC BY public copyright licence to any Author Accepted Manuscript version arising from this submission.

## Author contributions

C.W. and G.R. conceived the study. G.R. collected data. C.W. and G.R. performed the analyses and wrote and reviewed the manuscript.

## Competing interests

CW receives funding from GSK and MSD. The funders had no influence on this work, or its publication.

## Notes

### Summary of Updates

The previous version looked bad when opened in certain PDF readers (eg. Firefox, Okular). We hope that this version looks better for all.

## References

1. Burt, C. & Munafò, M. Has GWAS lost its status as a paragon of open science? PLOS Biol. 19, e3001242 (2021).

2. Thelwall, M. et al. Is useful research data usually shared? An investigation of genome-wide association study summary statistics. PLOS ONE 15, e0229578 (2020).

3. Buniello, A. et al. The NHGRI-EBI GWAS Catalog of published genome-wide association studies, targeted arrays and summary statistics 2019. Nucleic Acids Res. 47, D1005–D1012 (2019).

4. Wilkinson, M. D. et al. The FAIR Guiding Principles for scientific data management and stewardship. Sci. Data 3, 160018 (2016).

5. Bonomi, L., Huang, Y. & Ohno-Machado, L. Privacy challenges and research opportunities for genomic data sharing. Nat. Genet. 52, 646–654 (2020).

6. Wellcome Trust. Sharing Data from Large-scale Biological Research Projects: A System of Tripartite Responsibility. (2003).

7. Willer, C. J., Li, Y. & Abecasis, G. R. METAL: fast and efficient meta-analysis of genomewide association scans. Bioinformatics 26, 2190–2191 (2010).

8. Zhu, Z. et al. Causal associations between risk factors and common diseases inferred from GWAS summary data. Nat. Commun. 9, 224 (2018).

9. Bulik-Sullivan, B. K. et al. LD Score regression distinguishes confounding from polygenicity in genome-wide association studies. Nat. Genet. 47, 291–295 (2015).

10. Wallace, C. A more accurate method for colocalisation analysis allowing for multiple causal variants. PLOS Genet. 17, e1009440 (2021).

11. Privé, F., Arbel, J. & Vilhjálmsson, B. J. LDpred2: better, faster, stronger. Bioinformatics btaa1029 (2020) doi:10.1093/bioinformatics/btaa1029.

12. Hayhurst, J. et al. A community driven GWAS summary statistics standard. http://biorxiv.org/lookup/doi/10.1101/2022.07.15.500230 (2022) doi:10.1101/2022.07.15.500230.

13. Fecher, B., Friesike, S. & Hebing, M. What Drives Academic Data Sharing? PLOS ONE 10, e0118053 (2015).

14. Sayogo, D. S. & Pardo, T. A. Exploring the determinants of scientific data sharing:Understanding the motivation to publish research data. Gov. Inf. Q. 30, S19–S31 (2013).

15. Heeney, C., Hawkins, N., Vries, J. de, Boddington, P. & Kaye, J. Assessing the Privacy Risks of Data Sharing in Genomics. Public Health Genomics 14, 17–25 (2011).

16. Re-identifiability of genomic data and the GDPR. EMBO Rep. 20, e48316 (2019).

17. Homer, N. et al. Resolving Individuals Contributing Trace Amounts of DNA to Highly Complex Mixtures Using High-Density SNP Genotyping Microarrays. PLoS Genet. 4, e1000167 (2008).

18. Mongeon, P., Robinson-Garcia, N., Jeng, W. & Costas, R. Incorporating data sharing to the reward system of science: Linking DataCite records to authors in the Web of Science. Aslib J. Inf. Manag. 69, 545–556 (2017).

19. Hutchins, B. I., Davis, M. T., Meseroll, R. A. & Santangelo, G. M. Predicting translational progress in biomedical research. PLOS Biol. 17, e3000416 (2019).

20. Hutchins, B. I., Yuan, X., Anderson, J. M. & Santangelo, G. M. Relative Citation Ratio (RCR): A New Metric That Uses Citation Rates to Measure Influence at the Article Level. PLoS Biol. 14, e1002541 (2016).

21. MacArthur, J. A. L. et al. Workshop proceedings: GWAS summary statistics standards and sharing. Cell Genomics 1, 100004 (2021).

22. Christensen, G., Dafoe, A., Miguel, E., Moore, D. A. & Rose, A. K. A study of the impact of data sharing on article citations using journal policies as a natural experiment. PLOS ONE 14, e0225883 (2019).

23. Piwowar, H. A., Day, R. S. & Fridsma, D. B. Sharing Detailed Research Data Is Associated with Increased Citation Rate. PLOS ONE 2, e308 (2007).

24. Colavizza, G., Hrynaszkiewicz, I., Staden, I., Whitaker, K. & McGillivray, B. The citation advantage of linking publications to research data. PLOS ONE 15, e0230416 (2020).

25. Kidwell, M. C. et al. Badges to Acknowledge Open Practices: A Simple, Low-Cost, Effective Method for Increasing Transparency. PLOS Biol. 14, e1002456 (2016).

26. Piwowar, H. Value all research products. Nature 493, 159–159 (2013).

27. Travis Riddle. iCiteR: A Minimal Wrapper Around NIH’s ‘iCite’ API. (2019).

28. ICite, Hutchins, B. Ian & Santangelo, George. iCite Database Snapshots (NIH Open Citation Collection). (2022) doi:10.35092/YHJC.C.4586573.

29. Guerrero-Bote, V. P. & Moya-Anegón, F. A further step forward in measuring journals’ scientific prestige: The SJR2 indicator. J. Informetr. 6, 674–688 (2012).

30. Scimago Journal & Country Rank. https://www.scimagojr.com/ (2022).

31. R Core Team. R: A language and environment for statistical computing. (2021).

